# Bergamottin a CYP3A inhibitor found in grapefruit juice inhibits prostate cancer cell growth by downregulating androgen receptor signaling causing cell cycle block and apoptosis

**DOI:** 10.1101/2021.05.06.442999

**Authors:** Opalina Vetrichelvan, Priyatham Gorjala, Oscar Goodman, Ranjana Mitra

**Author notes:** Corresponding author (RM). Biomedical Sciences, College of Medicine, Roseman Uiversity of Health Sciences, 10530 Discovery Drive, Las Vegas, NV-89135.

## Abstract

Prostate cancer is the second leading cause of cancer related death in American men. Several therapies have been developed to treat advanced prostate cancer, but these therapies often have severe side effects. To improve the outcome with fewer side effects we focused on the furanocoumarin bergamottin, a natural product found in grapefruit juice and a potent CYP3A inhibitor. Our recent studies have shown that CYP3A5 inhibition can block androgen receptor (AR) signaling, critical for prostate cancer growth. We observed that bergamottin reduces prostate cancer (PC) cell growth by decreasing both total and nuclear AR (AR activation) reducing downstream AR signaling. Bergamottin’s role in reducing AR activation was confirmed by confocal microscopy studies and reduction in PSA levels. Further studies revealed that bergamottin promotes cell cycle block and accumulates G0/G1 cells. The cell cycle block was accompanied with reduction in cyclin D, cyclin B, CDK4, P-cdc2 (Y15) and P-wee1 (S642). We also observed that bergamottin triggers apoptosis in prostate cancer cell lines as evident by TUNEL staining and PARP cleavage. Our data suggest that bergamottin may be used as an adjunctive nutritional supplement to suppress prostate cancer growth and is of relevance to AA patients carrying wild type CYP3A5 often presenting aggressive disease.

## Introduction

Prostate cancer (PC) is the second leading cause of cancer death in US men: according to American Cancer Society approximately 34,130 men will die of prostate cancer in 2021. Anti-cancer treatment often has adverse effects due to its action within normal tissues. Given the initial slow growing nature of the prostate cancer and the front-line efficacy of androgen deprivation therapy (ADT), adjunctive therapy using dietary supplements with these ADT may provide us options to lower ADT dosage and reduce its adverse effect.

In this study we have tested the effect of bergamottin a natural occurring furanocoumarin found in grapefruit juice. Furanocoumarins are secondary metabolites produced in citrus fruits and they support plants defense against pathogens (1-3). The major furanocoumarins found in grapefruit are bergamottin, epoxybergamottin and 6’,7’ dihydroxybergamottin (4). Furanocoumarins have been reported to have anticancer activity and are known to activate multiple signaling pathways leading to cell cycle arrest, apoptosis and cell death (3, 5). Bergamottin has been studied for its anti-proliferative effect on different cancer cell types as a dietary supplement (6-8).

Bergamottin, the furanocoumarin compound used in our current study is also a strong inhibitor of CYP3A4/5 a P450 enzyme dominantly expressed in liver (9-12). Based on its crystal structure bergamottin shows a tendency towards π-π stacking which influences its interaction with the heme group of CYP3A4/5 (9). CYP3A5 is the main extrahepatic P450 form expressed in both normal prostate and in prostate cancer (13) hence bergamottin, a CYP3A5 can play an important role in treating prostate cancer. Androgen receptor (AR) is the driving force of prostate cancer growth, in our previous studies we have shown that CYP3A5 expressed in intratumoral prostate regulates AR expression regulating prostate cancer cell growth (14). We selected bergamottin for this study to monitor if it can be used as an CYP3A5 inhibitor to reduce AR activation and block prostate cancer growth.

Prostate cancer represents a major health disparity among African American (AA) men as new cases diagnosed are 60% higher in AAs as compared to Caucasian men (CA) (15, 16). One potential contributing fact to the health disparity is that AAs preferentially carry wild type CYP3A5 which constitutively promotes AR activation. CA men on the other hand carry the mutant form expressing very low amounts of CYP3A5 protein. As bergamottin is a CYP3A inhibitor its inhibitory effect can provide with a treatment strategy in AAs expressing high levels of CYP3A5 (17). In the current study to test the effect of bergamottin on prostate cancer growth we used two cell lines one of AA origin (MDAPCa2b) expressing wild type CYP3A5 and the other of CA origin (LNCaP) carrying mutant CYP3A5. Bergamottin has been shown to inhibit cancer cell growth by regulating multiple pathways in different cancer types, stat3 pathway, PARP cleavage and apoptosis, PI3/Akt survival pathway, cyclinD1/cell cycle block and VEGF/angiogenesis inhibition (3, 18, 19). So far it has not been tested for its ability to block AR activation which is very relevant in prostate cancer. In our recently published work we have shown how CYP3A5 inhibitors and inducers can alter AR activation more so in AAs expressing wild type CYP3A5 (20). Understanding the biological mechanisms how bergamottin regulates AR signaling in prostate cancer patients can help establish its mechanism based role in blocking prostate cancer growth especially in AAs expressing high CYP3A5 to reduce health disparity.

## Materials and Methods

### Cell lines, Drugs and antibodies

LNCaP, MDAPCa2b and RWPE1 cells were purchased from ATCC and maintained in RPMI (Invitrogen, Carlsbad, CA), F-12K medium (ATCC® 30-2004) and Keratinocyte SFM media (Invitrogen, Carlsbad, CA) respectively. Supplements were added as recommended by ATCC.

Antibodies against Androgen receptor (ab74272) were obtained from Abcam, (Cambridge, MA) anti-GAPDH (10R-G109A) was from Fitzgerald Industries (Acton, MA). Anti-α tubulin (2125S) and anti Lamin A/C (4C11), anti-PSA/KLK3 (D6B1) PARP (9542) and cell cycle regulation sampler kits (9932 and 9870) were obtained from Cell signaling technologies (Danvers, MA). The secondary antibodies (IR dye 680 and IR dye 800) were from LI-COR (Lincoln, NE). CYP3A inhibitor bergamottin (01338) was purchased from Sigma-Aldrich (St. Louis, MO).

### MTT assay

Growth of the cells was measured using MTT assay. Cells were plated (2000 cells/ 96 well or 13,000 cells/ 24 well) for 48 hours then treated for required time periods as mentioned and incubated with MTT (3-(4,5-Dimethylthiazol-2-yl)-2,5-Diphenyltetrazolium bromide) (4 mg/ml) for 2 h. at 37 °C. Cells were centrifuged at 2000 g for 10 min and the supernatant was discarded. The cell pellet was dissolved in 100/500 μl of DMSO. A plate reader was used to read the absorption at 540 nm. Experiments were performed in octuplet/quadruplet and repeated at least three times.

### Clonogenic assay

For clonogenic assay, cells were plated in 6 well plates (10,000 cells) for 48 h and then treated with indicated amounts of bergamottin. After 2 to 3 weeks, the colonies were fixed, stained (0.75% crystal violet, 50% ethanol and 1.75% formaldehyde) and counted. The assay was repeated three times.

### Western blotting

Cells were washed in phosphate buffer (PBS) and lysed in RIPA buffer (50 mM TRIS, 150 mM NaCl, 1% NP40, 0.5% sodium deoxycholate, 0.1% SDS, 2mM EDTA, 2mM EGTA) supplemented with protease and phosphatase inhibitors for total protein isolation. GAPDH was used as an internal control for total protein. Infrared fluorescent-labeled secondary antibodies were used for detection using Odyssey CLx. The bands were quantified using Odyssey software, which calculates pixel density and automatically takes an area adjacent to each band for background corrections.

### Cell fractionation

Nuclear and cytoplasmic cell fractionation was prepared using NE-PER Nuclear and cytoplasmic extraction kit from Thermo Scientific (Cat no.78833) and manufacturer’s instructions were followed. When the cells were treated with DHT, the media was changed to charcoal stripped serum media without phenol red (CSSM) 48 hours prior to the treatment. The cells were treated with bergamottin 72 hours after plating and 48 hours after changing the media to CSSM. Bergamottin and DHT were added at specified concentration and duration as indicated. Cells were washed in PBS once before cells were suspended in CER buffer. Protease, phosphatase inhibitors and EDTA was added prior to cell lysis. The pellet remaining after cytoplasmic isolation was washed twice with PBS. The pellet was suspended in NER buffer for nuclear fraction extraction according to guidelines, samples were stored at -80°C until further processing. Tubulin and Lamin were used as internal controls for cytoplasmic and nuclear fractions respectively.

### Confocal Microscopy

Cells were seeded into 35 mm Glass bottom dish (Cellvis catalog# D35C4-20-1.5-N). The cells were fixed in 4% paraformaldehyde for 20 minutes and permeabilized using permeabilizing buffer (0.2% Tween 20 in PBS) for 5 minutes. Cells were blocked using 10% goat serum diluted in permeabilizing buffer with 1% BSA for 15 minutes. Primary antibodies were diluted at 1:100 in staining buffer (1% BSA in PBS) and incubated for 2 hours at room temperature. Cells were washed three times (5 minutes each) in PBS. Secondary antibodies, TRITC-conjugated Donkey Anti-rabbit (711-025-152) was from Jackson immuno research, West Grove, PA was diluted at 1:100 in staining buffer and incubated for 60 minutes at room temperature. Cells were washed three times (5 minutes) in PBS and stained with 1 µg/mL DAPI (4’, 6-diamidino-2-phenylindole) in PBS for 5 minutes at room temperature. The cells were stored in PBS at 4°C until imaging was completed.

Cells were imaged using confocal laser scanning microscopy on a Nikon A1R using a galvano scanner and a 60× Apo-TIRF oil immersion objective. NIS Elements software form Nikon was used for recording the data.

### Cell cycle assay and analysis

Cell cycle analysis was performed after staining the cells with propidium iodide (PI). The cells were trypsinized washed with PBS and fixed in 70% ethanol (optional) and then stained with PI in nicoletti buffer (propidium iodide 50 μg/ml, 0.1% sodium citrate, 0.1% triton X-100, RNase 1 mg/ml, in DPBS). Cells were analyzed using C6 Accuri flow cytometer (Becton Dickinson, Mountainview, CA). FlowJo software was used to gate and remove the debris and aggregates from the analysis. Watson pragmatic method was used to assign percentage values to cells present in the different cell cycle stages.

### Apoptosis detection assay

To detect apoptosis in the cells after bergamottin treatment In Situ cell death detection kit from Sigma-Aldrich ((St. Louis, MO) was used. This apoptosis detection kit fluorescently labels the single and double-stranded DNA breaks caused due to apoptosis using terminal deoxynucleotidyl transferase (TUNEL-reaction). The cells were plated in 35 mm Glass bottom dish (Cellvis catalog# D35C4-20-1.5-N) treated with bergamottin and then stained with the kit reagents, the kit recommended protocol was followed. After the staining confocal microscope was used to procure images of the cells. The TUNEL assay preferentially labels apoptosis in comparison to necrosis and positive apoptosis is recognized when cells incorporate fluorescent labelled nucleotide (Ex 450 to 500nm and Em 515 to 565-green).

## Results

### Bergamottin blocks growth of prostate cancer cell lines

We tested the effects of bergamottin on prostate cancer cells using MTT and clonogenic assays. Since bergamottin is, a known CYP3A inhibitor we used two separate prostate cancer cell lines to test its effect differentially: LNCaP-of Caucasian origin cell line carrying mutant CYP3A5 and expressing low CYP3A5 and MDAPCa2b-of African American origin and carrying one wild type CYP3A5 and expressing higher levels of protein. MTT assay revealed that bergamottin reduced prostate cancer cell growth. The IC_50_ values for LNCaP and MDAPCa2b cells were observed to be 2.4 μM and 4 μM respectively (Fig 1A). Growth assay also revealed that bergamottin does not effect growth of RWPE1 cells (non transformed prostate epithelium) under the assay conditions. The growth inhibition effect of bergamottin was further confirmed using long term clonogenic assays. Bergamottin significantly reduced the number of colonies in both the cell lines LNCaP and MDAPCa2b at both 5μM and 10μM concentrations (Fig 1B).

**Fig 1:**
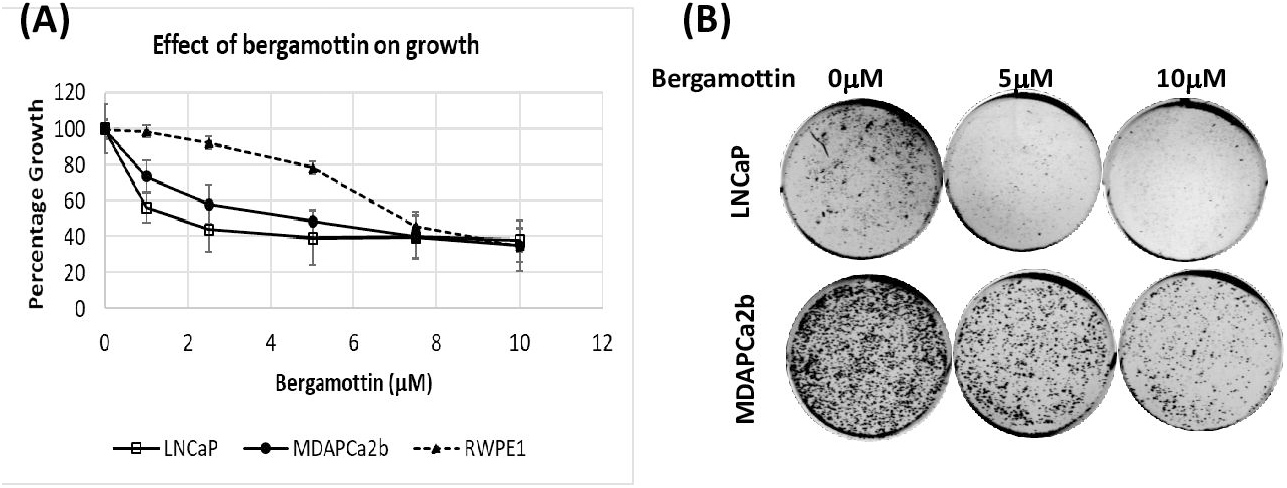
Bergamottin blocks prostate cancer cell growth. (A) Cells were treated with bergamottin for 72 hours; 48 hours after plating. MTT assay was performed as described in methods, the OD540 values were converted to relative percentage growth and plotted. (B) Clonogenic assay showing number of colonies formed after 2-3 weeks of bergamottin treatment in both the cell lines. The colonies were fixed and stained with crystal violet for visualization.

### Bergamottin reduces androgen expression, activation and downstream signaling

Androgen receptor is the driving force of prostate cancer growth. Since we observed that bergamottin blocked prostate cancer cell growth we tested its effect on AR expression and signaling. Bergamottin downregulated total AR levels in both LNCaP and MDAPCa2b cells by 50% (Fig 2A). DHT slightly increased the total AR expression in control and bergamottin treated LNCaP cells in only in control set in MDAPCa2b cells. Cell fractionation experiments with LNCaP cells showed that bergamottin inhibited nuclear translocation of AR consistent with CYP3A5 siRNA inhibition reported earlier: nuclear AR with and without DHT induction was lower in the bergamottin treated cells compared to control set (Fig 2B). The inhibition of nuclear translocation of AR was further confirmed in both the cell lines using confocal microscopy studies (Fig 2 C and D). In case of control (DMSO treated) cells we observe active AR migration to the nucleus with DHT induction, which is significantly reduced in bergamottin treated cells. Additionally, bergamottin also significantly reduced PSA levels in both cell lines which is a readout for AR downstream signaling (Fig 2E).

**Fig 2:**
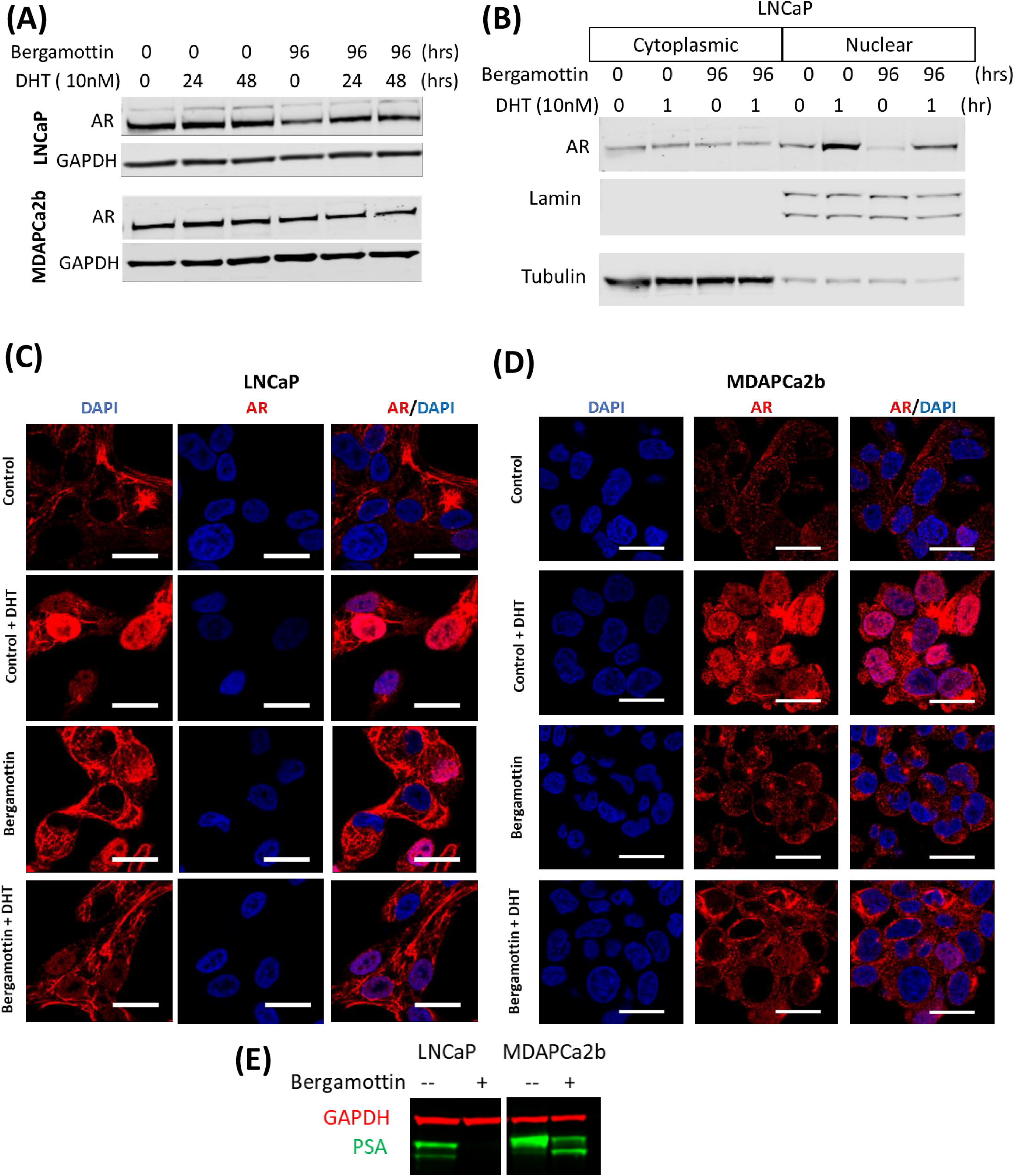
Bergamottin downregulated total AR expression, AR activation and downstream signaling: (A) Western blotting showed that bergamottin reduces total AR. GAPDH was used as an internal control for equal loading. Cells were grown in charcoal stripped serum and phenol red free media for 48 hours before DHT treatment. (B) Analysis of AR nuclear localization shows that bergamottin reduces AR in nuclear fraction with or without DHT induction. Lamin and tubulin have been used as internal controls. (C&D) Confocal microscopy showing effect of bergamottin on AR nuclear localization. Cells were plated in glass bottom plates and growth in phenol red free charcoal stripped serum media for 48 hours before DHT treatment (10nM). Size bar 20µm. (E) Western blotting showing reduction in PSA production with bergamottin treatment (10μM, 96 hours).

### Bergamottin induces cell cycle arrest

Since bergamottin reduces cell growth we tested its effect on cell cycle. Cell cycle analysis revealed accumulation of cells in the G1 phase and significant reduction of S-phase cells in the bergamottin treated set as compared to control (Fig 3). Both the analysis methods Watson pragmatic and Dean-Jett-Fox show accumulation of cells in the G0/G1 phase in both LNCaP and MDAPCa2b cells. Reduction in S phase (Table 1) was also observed which was more evident in LNCaP than MDAPCa2b cells.

**Fig 3:**
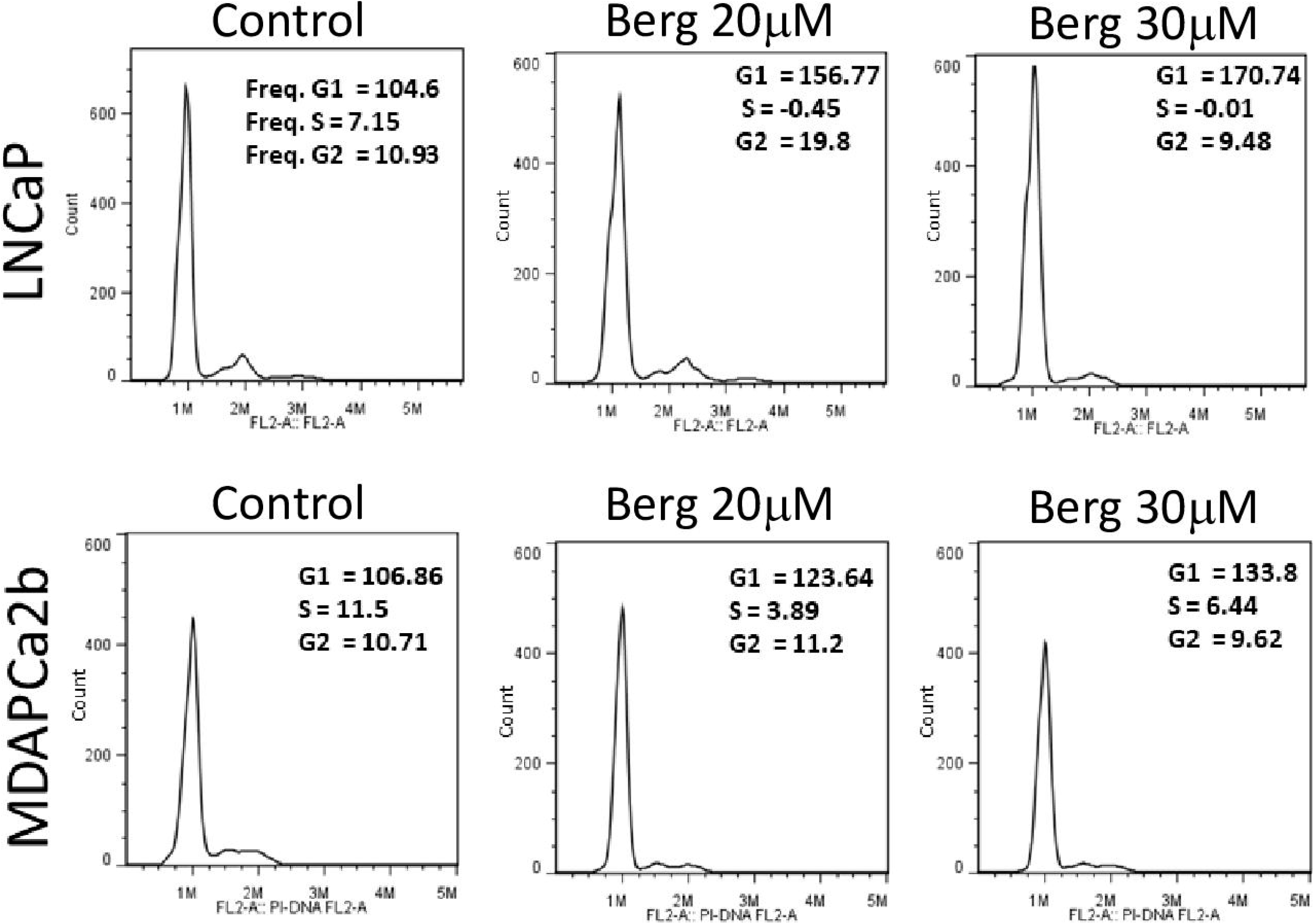
Bergamottin causes cell cycle block in prostate cancer cells. Cell cycle assay showing G0/G1, S and G2/M population in the control and bergamottin treated cells. The cell cycle analysis was performed as described in methods and the analysis was done using FlowJo software, a representative experiment is shown with Watson pragmatic analysis.

**Table 1:** Cell cycle analysis of bergamottin treated cells. Table showing increase in G0/G1 population and decrease in S phase cells with bergamottin treatment in both cell lines using both cell cycle analysis methods Dean-Jett-Fox and Watson pragmatic. The values represent average of three independent experiments, ± indicates SD values.

| LNCaP |  |  |  |  |  |  |  |
| --- | --- | --- | --- | --- | --- | --- | --- |
|  | Dean-Jett-Fox |  |  |  | Watson Pragmatic |  |  |
|  | G1 | S | G2 |  | G1 | S | G2 |
| DMSO | 75.7 ±4.2 | 9.7 ±4.8 | 11.8 ±0.7 | DMSO | 96.5 ±8.9 | 7.5 ±1.3 | 10.6 ±0.9 |
| Berg (20µM) | 86.7 ±2.3 | -9.8 ±3.0 | 15.3 ±1.0 | Berg (20µM) | 119.7 ±9.6 | 4.8 ±3.5 | 13.4 ±4.3 |
| Berg (30µM) | 96.2 ±3.6 | -8.4 ±4.5 | 7.7 ±0.9 | Berg (30µM) | 134.7 ±7.2 | 3.7 ±2.6 | 6.3 ±2.3 |

**Table1:** Cell cycle analysis of bergamottin treated cells.
| MDAPCa2b |  |  |  |  |  |  |  |
| --- | --- | --- | --- | --- | --- | --- | --- |
|  | Dean-Jett-Fox |  |  |  | Watson Pragmatic |  |  |
|  | G1 | S | G2 |  | G1 | S | G2 |
| DMSO | 76.8 ±1.9 | 11.4 ±1.7 | 10.5 ±1.3 | DMSO | 113.2 ±5.8 | 9.8 ±1.8 | 9.2 ±2.5 |
| Berg (20µM) | 84.9 ±0.4 | 6.7 ±0.7 | 9.6 ±0.5 | Berg (20µM) | 122.5 ±3.7 | 7.1 ±1.7 | 9.6 ±1.8 |
| Berg (30µM) | 88.4 ±3.6 | 3.3 ±3.4 | 6.8 ±3.2 | Berg (30µM) | 135.5 ±3.0 | 5.6 ±0.5 | 8.4 ±1.1 |

### Western blot analysis showing differential expression of cell cycle regulating proteins after bergamottin treatment

Cell cycle arrest occurs due to block at one of the four check points: G1/S check point, G2/M check point, SAC check point, intra S check point and restriction point (reversable). All these check points are regulated by cyclins and CDKs, we tested the effect of bergamottin on these cell cycle regulating proteins. We first tested effect of bergamottin on cyclins and CDKs and then on proteins regulating cyclins and CDKs. Our western blot analysis reveal reduction of Cyclins B1, D1, D3 and E in LNCaP cells and reduction of Cyclins A, B1 and D3 in MDAPCa2b cells (Fig 4A, Table 2). We observed reduction of CDK4 and CDK6 in LNCaP and only CDK4 in MDAPCa2b cells. The fold changes after GAPDH calibration are shown in Table 2.

**Fig 4:**
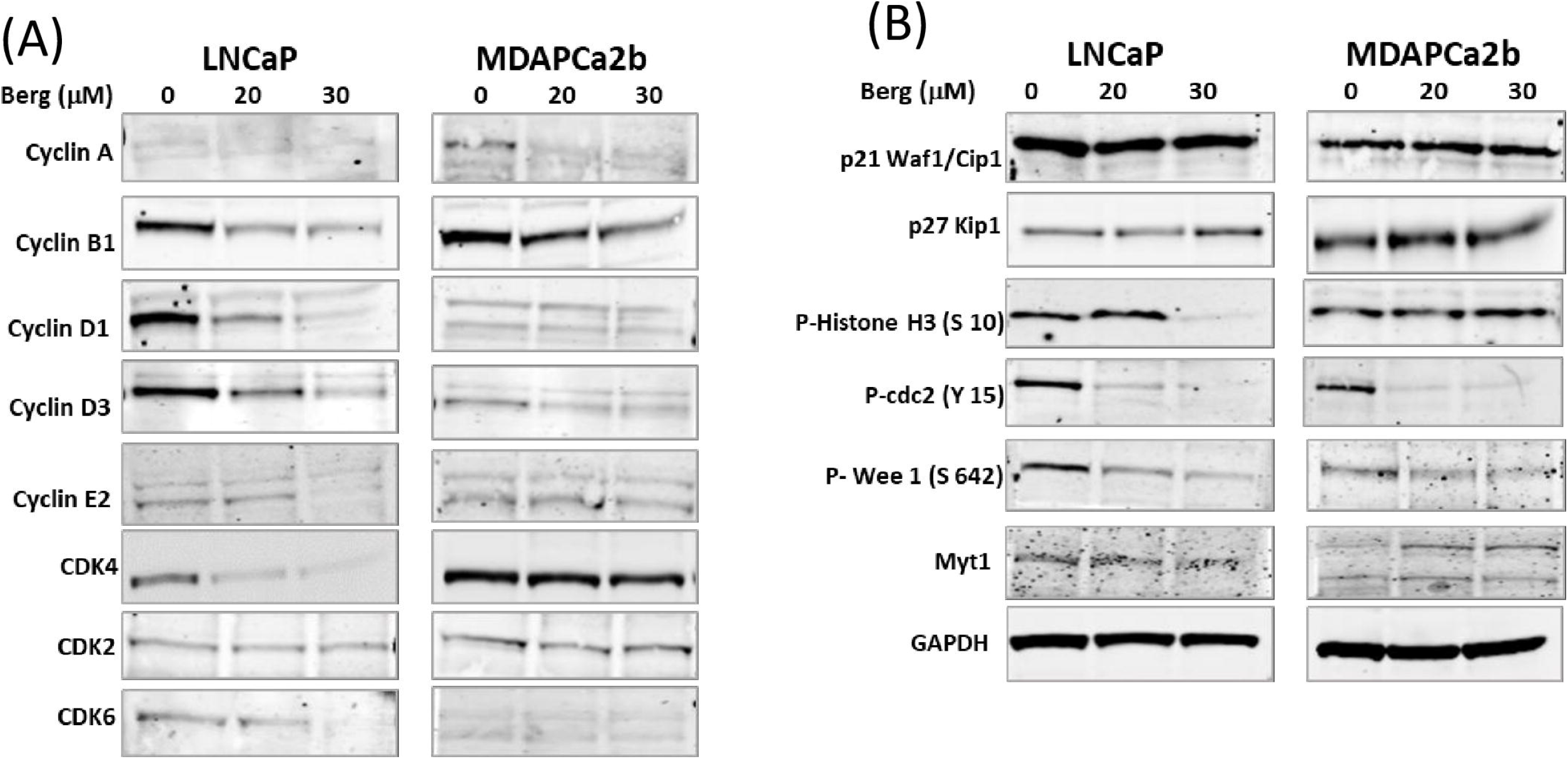
Differential regulation of cell cycle regulating protein after bergamottin treatment. (A &B) Western blot analysis of control and bergamottin treated cells depicting changes in cell cycle regulating proteins. GAPDH has been used as a loading control. The experiment was repeated three times, a representative replicate is shown.

**Table 2:** Fold changes in cell cycle regulating proteins after bergamottin treatment.

|  | LNCAP |  |  | MDAPCa2b |  |  |
| --- | --- | --- | --- | --- | --- | --- |
| | Bergamottin (μM) | | | Bergamottin (μM) | | |
|  | 0 | 20 | 30 | 0 | 20 | 30 |
| Cyclin A | 100 | 80 | 55 | 100 | 86 | 58 |
| Cyclin B1 | 100 | 50 | 44 | 100 | 52 | 42 |
| Cyclin D1 | 100 | 43 | 12 | 100 | 106 | 110 |
| Cyclin D3 | 100 | 68 | 44 | 100 | 55 | 49 |
| Cyclin E | 100 | 109 | 53 | 100 | 120 | 87 |
| CDK2 | 100 | 62 | 67 | 100 | 63 | 77 |
| CDK4 | 100 | 46 | 30 | 100 | 90 | 81 |
| CDK6 | 100 | 60 | 12 | 100 | 108 | 98 |
| p21 | 100 | 83 | 102 | 100 | 139 | 126 |
| p27 | 100 | 112 | 183 | 100 | 158 | 143 |
| P-histone H3 | 100 | 294 | 35 | 100 | 130 | 152 |
| **P-cdc2** | 100 | 18 | 5 | 100 | 14 | 15 |
| **P-Wee1** | 100 | 63 | 37 | 100 | 80 | 87 |
| **Myt 1** | 100 | 103 | 82 | 100 | 259 | 451 |

Additionally, we observed increase in p27kip1 expression in both cell lines and no changes were observed in p21 levels. Decreased phosphorylation of cdc-2 (Y15) and wee-1 (S642) were also observed in both cell lines. Changes in phosphorylation of Histone H3 (S10) and Myt1 levels differed between the two cell lines with bergamottin treatment.

Table represents normalized fold changes after GAPDH calibration and corresponds to the replicate shown in Fig 4.

### Bergamottin causes cell death by triggering apoptosis in the cells

Since we observed cell cycle blockade and growth inhibition with bergamottin treatment, we wanted to test if it is triggering apoptosis. To assess apoptosis we performed tunnel assays with both the cell lines (Fig 5 A & B). The tunnel assay indicates DNA breaks (green fluorescence) due to apoptosis with bergamottin treatment in both cell lines. Additionally, both the cell lines show PARP cleavage, another indicator of apoptosis (Fig 5C).

**Fig 5:**
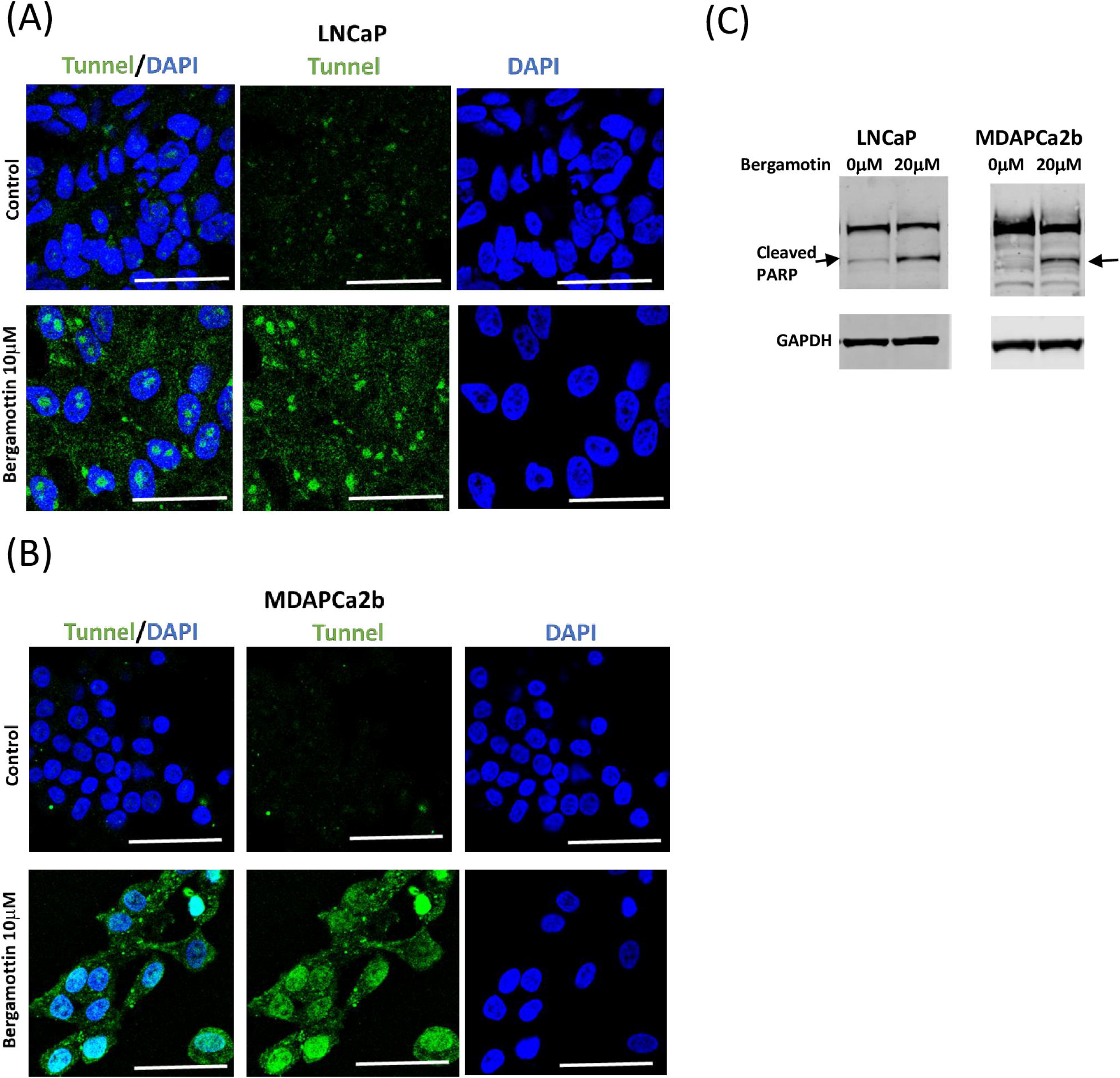
Bergamottin causes apoptosis in LNCaP and MDAPCa2b cells. (A & B) LNCaP and MDAPCa2b cells were treated with bergamottin (10µM) for 4 days. Cells were then fixed and stained for tunnel assay reagents. Size bar 50µm. DAPI was used to stain the nucleus. (C) Western blot analysis showing cleaved PARP after bergamottin treatment, GAPDH was used as an internal control.

## Discussion

Our results indicates that bergamottin inhibits growth in both LNCaP and MDAPCa2b cells at significantly low concentrations (IC_50_ values, 2.4 and 4µM respectively). Previously bergamottin has been shown to reduce growth of several other cancer cell types but the IC_50_ were reported in the range of 30-100µM (18, 19, 21-29). The clonogenic assays also show significant loss of growth at 5 and 10 µM concentrations. This may be because both the cells express CYP3A5 active protein and bergamottin is a potent CYP3A5 inhibitor. The lower IC_50_ values for LNCaP may be attributed to lower levels of CYP3A5 protein due to *3/*3 mutation present as compared to MDAPCa2b which carries one copy of wild type of CYP3A5 (*1/*3) (20). The other major difference between the previously reported results in the DU145 AR negative prostate cancer cell line (18) suggests bergamottin’s action is CYP3A5 - AR signaling dependent (14). Bergamottin does not effect RWPE1 (non-transformed prostate epithelium) cells at the tested concentration which is consistent with the earlier reported data (18).

We have reported earlier that CYP3A5 siRNA blocks AR nuclear translocation (14) and bergamottin which is a CYP3A inhibitor (6, 10, 30) also shows reduced nuclear AR consistent with our earlier results. Although primarily bergamottin is a CYP3A4 inhibitor it also inhibits CYP3A5, CYP2B6 and CYP1A1 (9, 11, 12, 31, 32). Since CYP3A5 is the major isoform expressed in normal prostate and prostate tumor the effects observed in our experiments is primarily due to inhibition of CYP3A5 in the tested prostate cancer cell lines (13, 33). Additionally, bergamottin also lowers total AR expression which may be further contributing to growth inhibition, as AR is known to be the driver of prostate cancer growth. We observed loss of PSA in the bergamottin treated cells, with PSA levels representing a downstream readout of AR signaling (34). This observation can be useful in patients with high AR and CYP3A5 expression such as African Americans, who preferentially carry wild type CYP3A5 and express higher AR, accompanied by relatively aggressive disease. Our experiments show accumulation of cells in the G0/G1 phase and depletion of S phase cells with bergamottin treatment. Accumulation of G0/G1 cells can occur at the restriction point or at the G1/S check point (35). The restriction point is regulated by cyclin D and CDK4,6 (36). Since we observed reduction in Cyclin D1, D3 and CDK 4,6 in LNCaP cells and Cyclin D3 and CDK4 in MDAPCa2b cells (Figure 4A, Table 2) it shows that the block is at the restriction point in both the cell lines. Additionally, we observed increase in p27kip1 which indicates increased population of quiescent cells (G0). Cyclin E and CDK2 are known to regulate G1/S check point which also leads to accumulation of G0/G1 cells, which is consistent with larger block observed in LNCaP cells which show downregulation of cyclin E2. Additionally, in LNCaP both Cyclin D1 and D3 are down regulated and in MDAPCa2b only Cyclin D3 is, which explains more pronounced accumulation of G1 cells in LNCaP as compared to MDAPCa2b. We also observed slight reduction in Cyclin A in bergamottin treated MDAPCa2b cells, cyclin A levels increase in S phase and is associated with intra S check point which can explain the why the loss of S phase cells is not evident in MDAPCa2b cells as compared to LNCaP. Both the cell lines showed reduced expression of cyclin B after bergamottin treatment which is associated with G2/M check point. Although we observe changes in the cyclin B but our cell cycle analysis does not show any increase in G2/M cell population. We observed decreased phosphorylation of cdc-2 (Y15), phosphorylation at Y15 is known to block mitotic entry, it is possible that the reduction in cyclin B and cdc-2 dephosphorylation cancel each other and we do not see any accumulation of G2/M cells (37). We observed less phosphorylation of wee-1 (S642) which is a Y15 kinase consistent with the loss of phosphorylation of cdc-2. There is loss of phosphorylation of Histone H (S10) in the 30uM bergamottin treated LNCaP cells which is essential for chromosomal condensation for segregation during mitosis. Bergamottin has been shown to increase sub-G1 population in myeloma cells and cause G2/M block in lung cancer cells at higher concentrations (18, 19)

Bergamottin induces apoptosis in both prostate cancer cell lines, evidenced by tunnel assay stanning of late apoptotic cells by labeling DNA breaks and nicks by Terminal deoxynucleotidyl transferase, which incorporates fluorescein UTP at 3’-OH ends in a template independent manner. Tunnel assay preferentially labels DNA strand breaks generated during apoptosis which allows discrimination of apoptosis from necrosis. Both the cell lines show apoptosis after bergamottin treatment, although MDAPCa2b cells have more fluorescein UTP incorporation than in LNCaP cells. In addition, we also observed PARP-cleavage which is an hallmark of apoptosis (38) and the irreversible binding of small cleaved PARP protein prevents DNA repair. Several anticancer drugs target androgen receptor signaling to prevent tumor growth, our observation that bergamottin a natural furanocoumarin found in grapefruit juice also blocks the androgen receptor expression and activation is very significant as it can be used as a dietary supplement to prevent prostate cancer growth. Previously we have investigated how CYP3A5 inhibitors and inducers can modify AR signaling and effect the efficacy of commonly prescribed concomitant medications (20). CYP3A4/5 are hepatic enzymes and are involved in processing several commonly prescribed drugs, we do have to be cautious of drug-drug interactions while using bergamottin as an anticancer agent as it is a strong CYP3A inhibitor. Use of bergamottin on the other hand can be very significant for African Americans who often present aggressive disease have high AR expression (39, 40) and often carry wild type CYP3A5 (41-43). Both, the relative incidence and mortality rate are much higher (1.8 and 2.2 times more respectively) in AAs as compared to Caucasian Americans (CA) (16). Since bergamottin is a natural CYP3A5 inhibitor combination with different other cancer drugs may provide an opportunity to increase efficacy of other cancer drugs especially in AAs expressing high levels of CYP3A5.

## Conclusions

Bergamottin a potent CYP3A inhibitor blocks prostate cancer cell growth by inhibiting AR expression and activation leading to cell cycle block and apoptosis. These observations suggest the potential use of bergamottin as a dietary supplement for prostate cancer prevention and treatment.

## Supporting information

**S1 Fig: Full blot of western gels shown in Fig 2A (A), Fig 2B (B) and Fig 2E (C)**. The dotted lines show places were the blots were cut to treat with separate antibodies. In (C) the IR dye 700 (red, GAPDH) and IR dye 800 (green, PSA) are shown a two different intensities to show clarity representative of the original figure 1E.

**S2 Fig: Full blot of western gels shown in figure 4A and 4B. (A)**- Blots with LNCaP extracts, **(B)**- Blots with MDAPCa2b extracts. The dotted lines show places were the blots were cut to treat with separate antibodies. During loading sometime, the markers are not next to the samples, in those cases the representative marker is placed next to the blot.

**S3 Fig: Full blot of western gels shown in Fig 5**.

