## Supplementary figures and images for "Bergamottin a CYP3A inhibitor found in grapefruit juice inhibits prostate cancer cell growth by downregulating androgen receptor signaling causing cell cycle block and apoptosis"

### Supplemental Fig 1

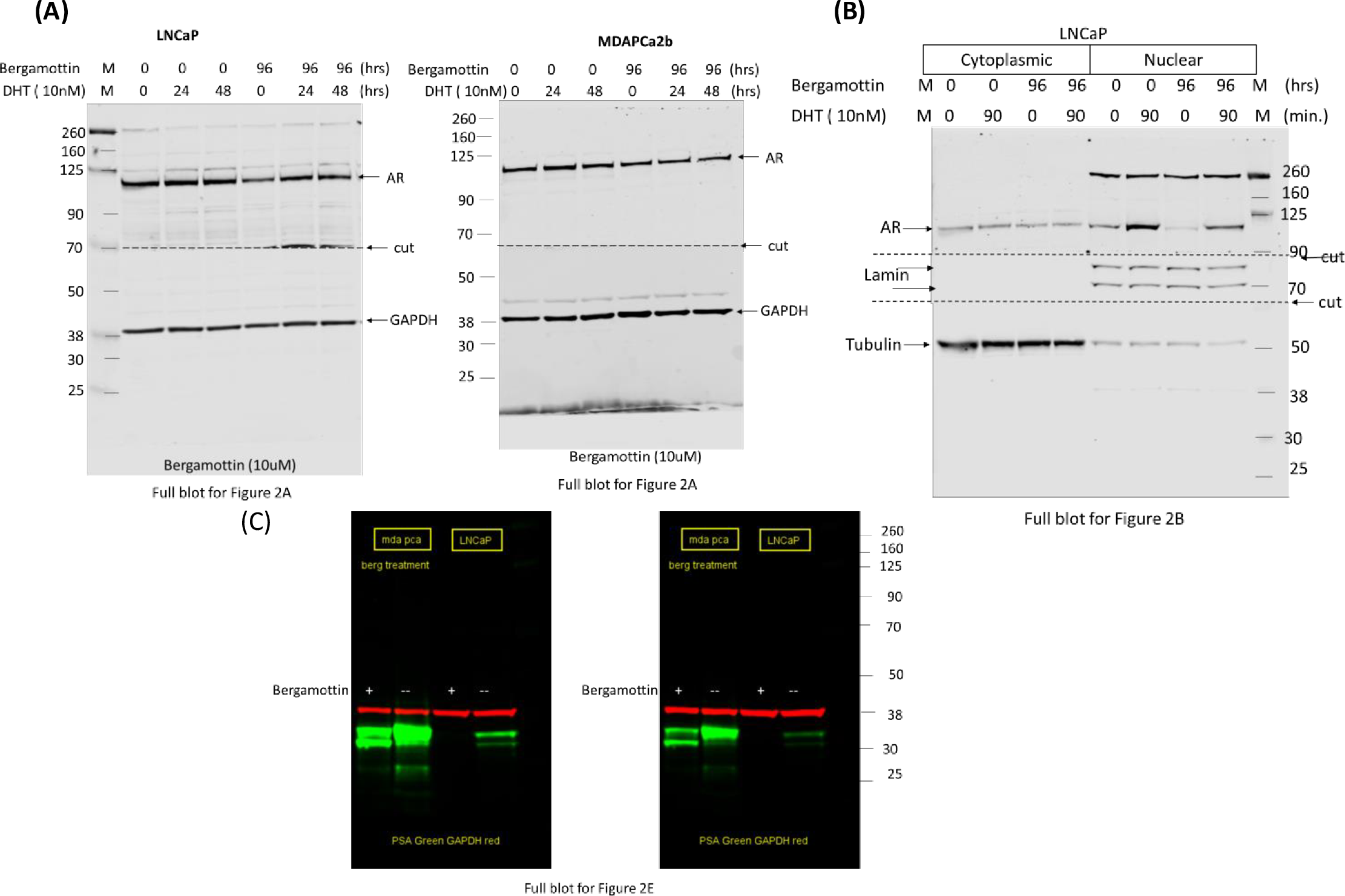

### Supplemental Fig 2

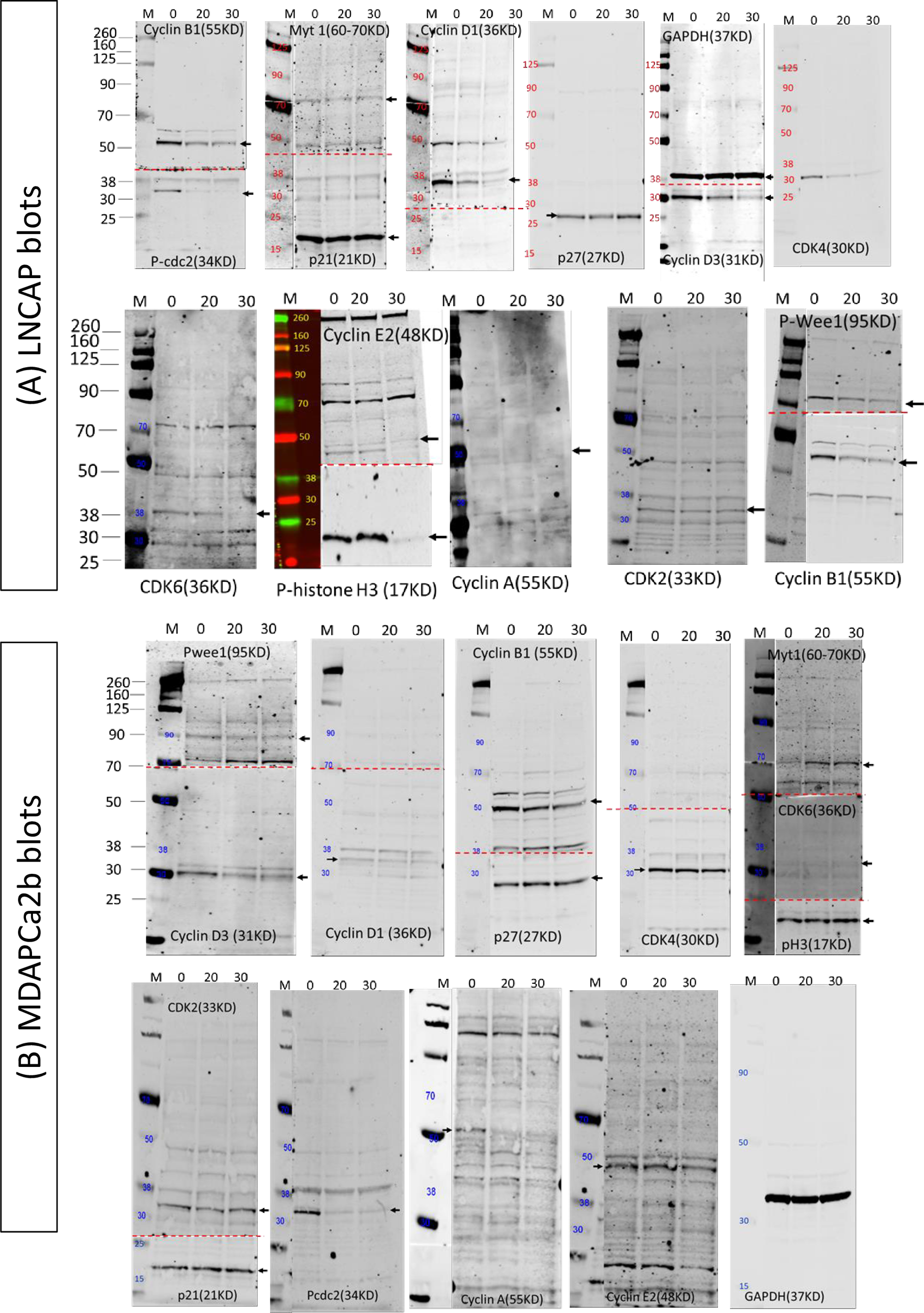

### Supplemental Fig 3

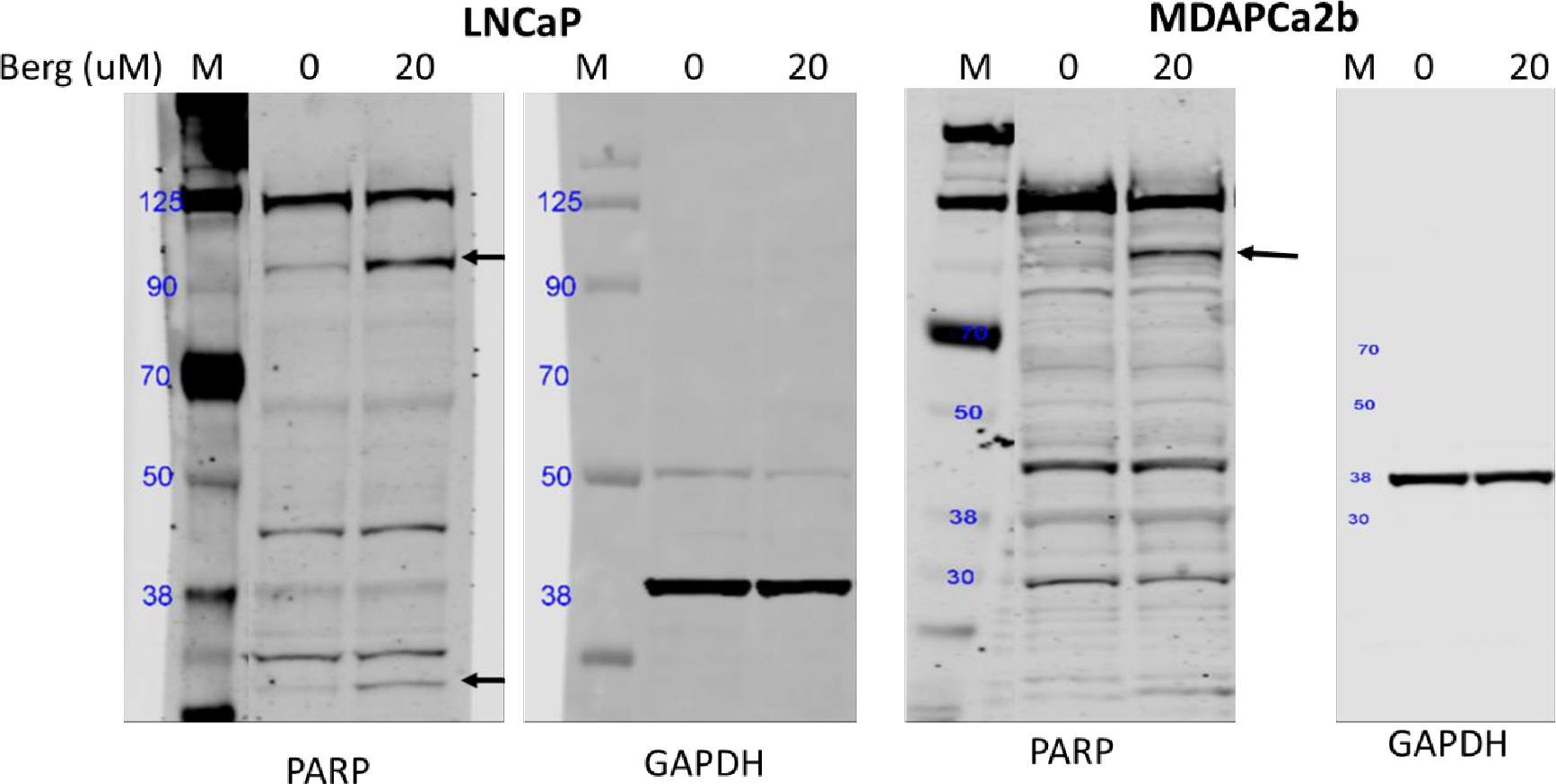
